# Region-specific cortico-striatal transcriptomic remodeling following early postnatal dopaminergic disturbance

**DOI:** 10.64898/2026.05.20.726444

**Authors:** Miyuki Doi, Stefano Berto, Shoichi Shimada, Noriyoshi Usui

## Abstract

Dopamine signaling plays critical roles in postnatal brain development, yet the molecular consequences of early dopaminergic disturbance remain incompletely understood. Here, we investigated transcriptomic alterations in the prefrontal cortex (PFC) and striatum (STR) of mice subjected to early postnatal dopaminergic disturbance by 6-hydroxydopamine (6-OHDA) treatment. Using bulk RNA sequencing (RNA-seq) and weighted gene co-expression network analysis (WGCNA), we identified 369 differentially expressed genes (DEGs) in the PFC, 493 DEGs in the STR, and 32 co-expression modules with region-specific expression patterns. Functional enrichment analyses showed that PFC DEGs were associated with cortical development, plasma membrane signaling, and transcriptional regulation, whereas STR DEGs were enriched for striatal development, locomotion, extracellular matrix organization, and amphetamine response. Co-expression network analysis further identified module-specific enrichments related to developmental, synaptic, metabolic, immune-related, and transcriptional programs. DEG sets from both regions also overlapped with genes implicated in attention-deficit/hyperactivity disorder (ADHD) and other neuropsychiatric disorders. Together, these findings reveal region-specific cortico-striatal transcriptomic remodeling following early postnatal dopaminergic disturbance and identify molecular programs that may link developmental dopaminergic perturbation to later behavioral phenotypes.

**Highlights:**

- Early dopaminergic disturbance reshapes cortico-striatal transcriptomes
- PFC changes were linked to developmental and transcriptional programs
- STR changes were linked to locomotion and extracellular matrix programs
- Network analysis revealed region-specific developmental and synaptic programs

## Introduction

Dopamine signaling is a key regulator of brain development and function, influencing neuronal maturation, circuit formation, synaptic activity, and behavior. Disruption of dopaminergic signaling during early life can have lasting effects on neural development and later behavioral outcomes. ADHD is a common neurodevelopmental disorder characterized by attention deficits, hyperactivity, and impulsivity^1,2^. Genetic, imaging, and genomic studies have implicated dopamine-related and neurodevelopmental pathways, as well as cortico-striatal circuit abnormalities, in ADHD^1,3-9^. Together, these findings suggest that altered dopaminergic and developmental processes may contribute to later behavioral phenotypes.

The PFC and STR are major components of cortico-striatal circuits that regulate attention, impulse control, motivation, and locomotor activity^1,4,8,9^. Dopamine plays essential roles in the maturation and function of these circuits, and perturbation of dopaminergic signaling during development may alter region-specific molecular programs with long-term consequences. Defining the molecular responses of these regions to early dopaminergic disturbance may therefore provide insight into developmental mechanisms linked to later behavioral abnormalities.

Animal models have provided important insights into how developmental dopaminergic perturbations influence later behavioral phenotypes. Genetic, spontaneous, and pharmacological models display phenotypes such as hyperactivity, impulsivity, and attention deficits^10-14^. Among non-genetic approaches, early postnatal treatment with 6-OHDA has been widely used as an experimental model of dopaminergic disturbance during brain development^15,16^. Because 6-OHDA damages both dopaminergic and noradrenergic neurons, desipramine is co-administered to protect noradrenergic neurons and thereby enrich for dopaminergic depletion^10-12,15-19^. This model exhibits behavioral alterations, including hyperactivity, impulsivity, and attention deficits, and some of these phenotypes are ameliorated by ADHD medications^10-12,15-19^. In our previous studies, mice treated with 6-OHDA during early postnatal development exhibited axon initial segment (AIS) shortening in the PFC and primary somatosensory barrel cortex, suggesting altered neuronal activity transmission and cortical development^15,16^.

However, the molecular consequences of early postnatal dopaminergic disturbance in cortico-striatal regions remain incompletely understood. In this study, we investigated transcriptomic changes in the PFC and STR at an early postnatal stage in mice treated with 6-OHDA. Using bulk RNA-seq and WGCNA, we identified region-specific DEGs and co-expression networks associated with developmental, synaptic, metabolic, and immune-related processes. These findings define a region-specific cortico-striatal transcriptomic landscape following early postnatal dopaminergic disturbance and provide a molecular framework for understanding molecular pathways linked to later behavioral abnormalities.

## Results

### Early postnatal dopaminergic disturbance induces hyperactivity and impulsivity-like behavior

To investigate the molecular consequences of early postnatal dopaminergic disturbance, we employed a postnatal 6-OHDA treatment model^15,16^ (Figure 1A). Following 6-OHDA administration, TH immunoreactivity was reduced in fibers projecting from dopaminergic neurons in the ventral tegmental area to the STR, indicating impaired dopaminergic innervation (Figure 1B).

**Figure 1.**
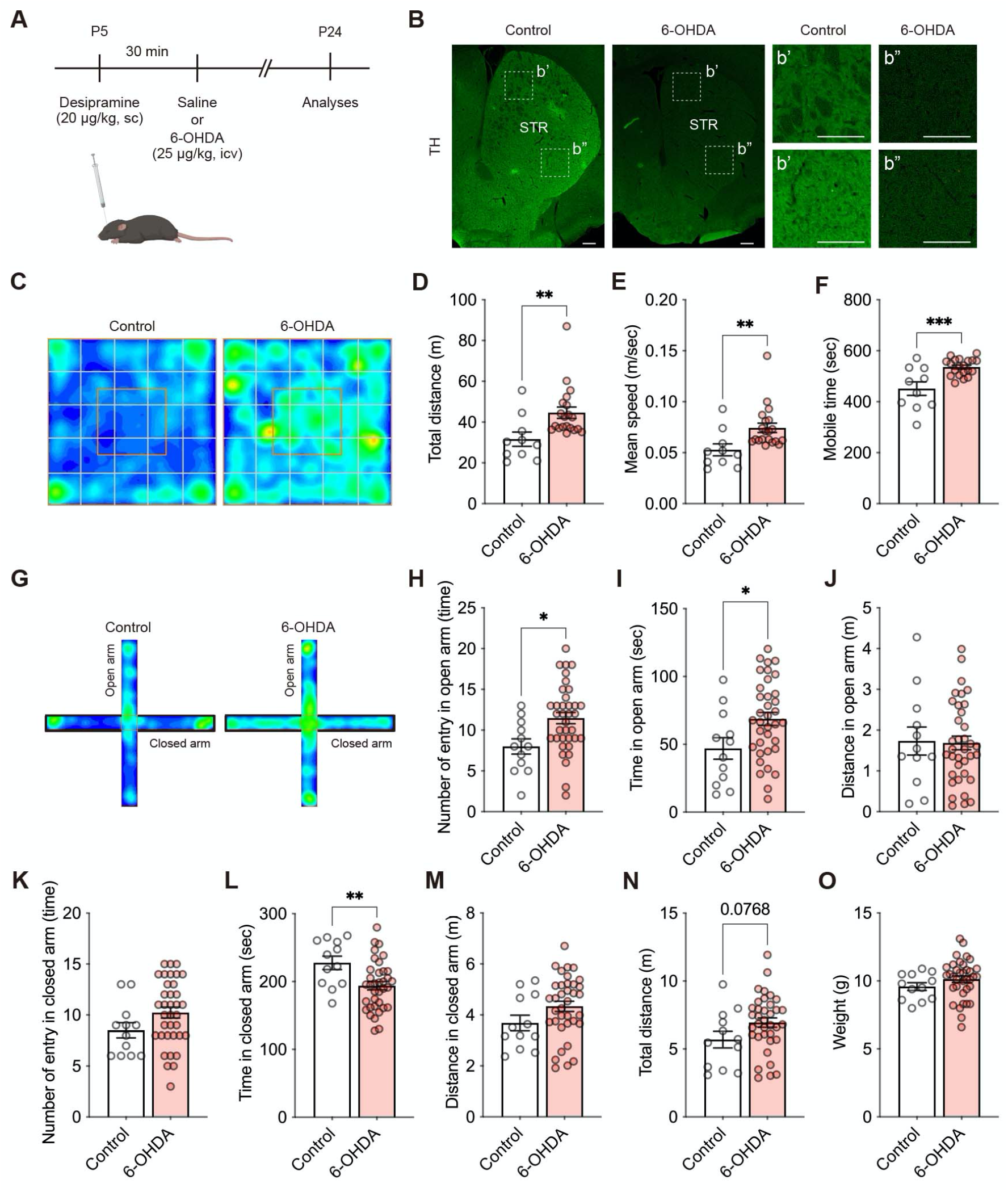
Early postnatal 6-OHDA treatment induces hyperactivity and impulsivity-like behavior in mice. (**A**) Experimental time course. Created in BioRender. (**B**) Representative fluorescent images of tyrosine hydroxylase (TH)-positive fibers from dopaminergic neurons in the striatum (STR) of mice at P24. **B’** and **B”** show enlarged views of **B**. TH-immunoreactivity was decreased in the STR of 6-OHDA-treated dopaminergic disturbance model mice. (**C**) Representative heatmaps of exploratory activity in the open field test. (**D-F**) Quantification of total distance travelled (**D**), mean speed (**E**), and mobile time (**F**) in the open field test. 6-OHDA-treated mice exhibited hyperactivity. (**G**) Representative heatmaps of mouse exploratory activity in the elevated plus maze test. (**H-J**) Quantification of the number of entries (**H**), time spent (**I**), and distance traveled (**J**) in the open arm. (**K-M**) Quantification of the number of entries (**K**), time spent (**L**), and distance traveled (**M**) in the closed arm. (**N**) Quantification of total distance in the elevated plus maze test. 6-OHDA-treated mice also exhibited impulsivity-like behavior. (**O**) Quantification of body weight. Data are represented as means (±SEM). Asterisks indicate ***P<0.001, **P<0.01, *P<0.05, unpaired *t*-test, n=10-20/condition for the open field test, n=12-36/condition for the elevated plus maze test, n=12-36/condition for body weight. Scale bar: 200 μm.

To characterize the behavioral effects of early postnatal dopaminergic disturbance, we performed open field and elevated plus maze tests. 6-OHDA-treated mice showed increased total distance traveled, mean speed, and mobile time compared to controls in the open field test (Figure 1C-F). In the elevated plus maze, 6-OHDA-treated mice displayed increased activity in the open arm area and a greater number of entries into this area (Figure 1G-J), whereas time spent in the closed arms was reduced (Figure 1G and K-M). A trend toward increased total distance traveled was also observed in the elevated plus maze test (Figure 1N). No significant difference in body weight was detected between groups (Figure 1O). These results indicate that early postnatal dopaminergic disturbance induces hyperactivity and impulsivity-like behavior.

### Region-specific transcriptomic alterations in the PFC following early postnatal dopaminergic disturbance

Next, we analyzed transcriptomic changes in the PFC and STR of 6-OHDA-treated mice at P24 using bulk RNA-seq. In the PFC transcriptome, transcriptomic profiles showed clear separation between groups (Figure 2A). We identified 369 DEGs in the PFC, including *Igfn1, Rims3, Etnppl, Cbs, Kmt2a, Ryr2, Mtss2, Htra1, Marcksl1, and Tubb2a* (Figure 2B, C and Table S1). GO analysis showed that PFC-DEGs were enriched in anatomical structure development, multicellular organism development, and regulation of developmental process (Figure 2D, Table S2, and Figure S2).

**Figure 2.**
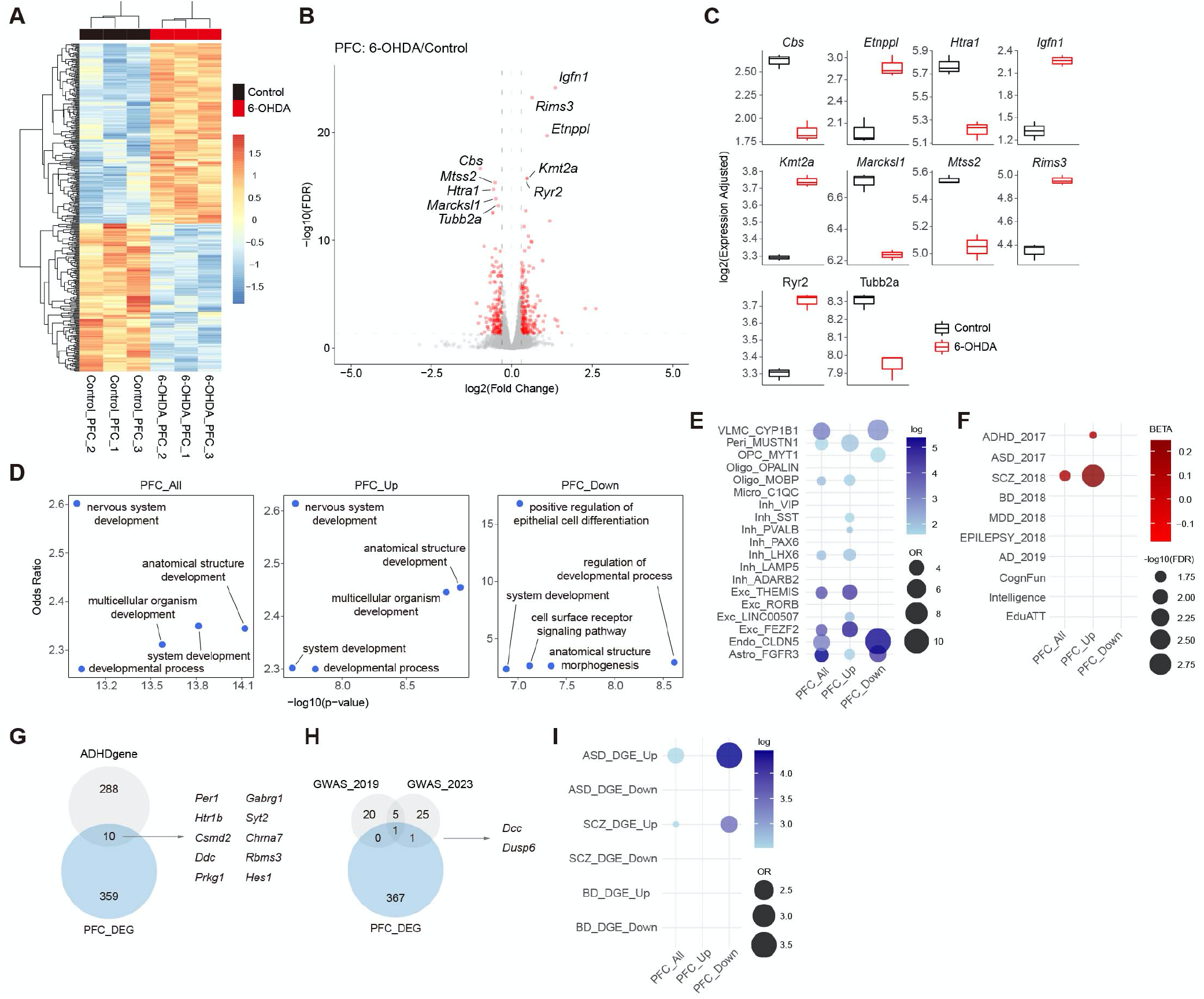
Transcriptomic remodeling in the PFC of 6-OHDA-treated mice. (**A**) Heatmap of differentially expressed genes (DEGs) in the mouse prefrontal cortex (PFC) at P24. (**B**) Volcano plot of PFC-DEGs. (**C**) Bar plots showing the top 10 PFC-DEGs. (**D**) Gene Ontology (GO) analysis of biological process (BP) terms for PFC-DEGs in all, upregulated, and downregulated genes, respectively. (**E**) Dot plots showing enrichment of PFC-DEGs across cell types. *VLMC: vascular lepotomeningeal cells, Peri: pericytes, OPC: oligodendrocyte precursor cells, Oligo: oligodendrocytes, Micro: microgia, Inh: inhibitory neurons, Exc: excitatory neurons, Endo: endothelial cells, Astro: astrocytes*. (**F**) Dot plots showing MAGMA beta values. *ADHD: attention deficit hyperactivity disorder, ASD: autism spectrum disorder, SCZ: schizophrenia, BD: bipolar disorder, MDD: major depressive disorder, Epilepsy: epilepsy, AD: Alzheimer’s disease, CognFunc: cognitive functions, Intelligence: intelligence, EduATT: educational attainment*. (**G**) Venn diagram showing overlap between PFC-DEGs and ADHDgene. Representative overlapping genes, including *CSMD2* and *DDC*, are highlighted. (**H**) Venn diagram showing overlap between PFC-DEGs and GWAS-associated genes. Representative overlapping genes, including *DUSP6, DCC*, and *FOXP2*, are highlighted. (**I**) Dot plots showing enrichment of PFC-DEGs in disorder-specific human transcriptomic datasets.

Cell type enrichment analysis revealed that PFC DEGs were enriched in vascular leptomeningeal cells (VLMCs), pericytes, endothelial cells, inhibitory and excitatory neurons, oligodendrocytes, and astrocytes (Figure 2E). GWAS enrichment analysis showed enrichment for genes associated with ADHD and schizophrenia (SCZ) (Figure 2F). In addition, 10 PFC DEGs (*Per1, Htr1b, Csmd2, Ddc, Prkg1, Gabrg1, Syt2, Chrna7, Rbms3*, and *Hes1*) overlapped with ADHDgenes^20^ (Figure 2G), and 7 PFC DEGs (*Kdm4a, Ptprf, Foxp2, Sorcs3, Poc1b, Dcc, Dusp6*) overlapped with two GWAS datasets^5,6^ (Figure 2H). PFC DEGs were also enriched among upregulated human DEGs associated with autism spectrum disorders (ASD) and SCZ (Figure 2I). These findings indicate that early postnatal dopaminergic disturbance induces region-specific transcriptomic remodeling in the PFC linked to developmental, neuronal, glial, and vascular-associated programs.

### Region-specific transcriptomic alterations in the STR following early postnatal dopaminergic disturbance

In the STR, transcriptomic profiles also showed clear separation between groups, similar to the PFC (Figure 3A). We identified 493 DEGs in the STR, including *Fmod, Fabp7, Slc22a6, Traf3, Thbs4, Smox, Col3a1, Serpinh1, H1f2*, and *Fn1* (Figure 3B, C, and Table S3). Among these, 93 DEGs were shared between the PFC and STR (Figure S3). GO analysis showed that STR DEGs were enriched in STR development, locomotion, and anatomical structure morphogenesis (Figure 3D, Table S4, and Figure 2). Notably, upregulated STR DEGs were particularly associated with STR development, locomotory behaviors, and response to amphetamine (Figure 3D, Table S4, and Figure 2).

**Figure 3.**
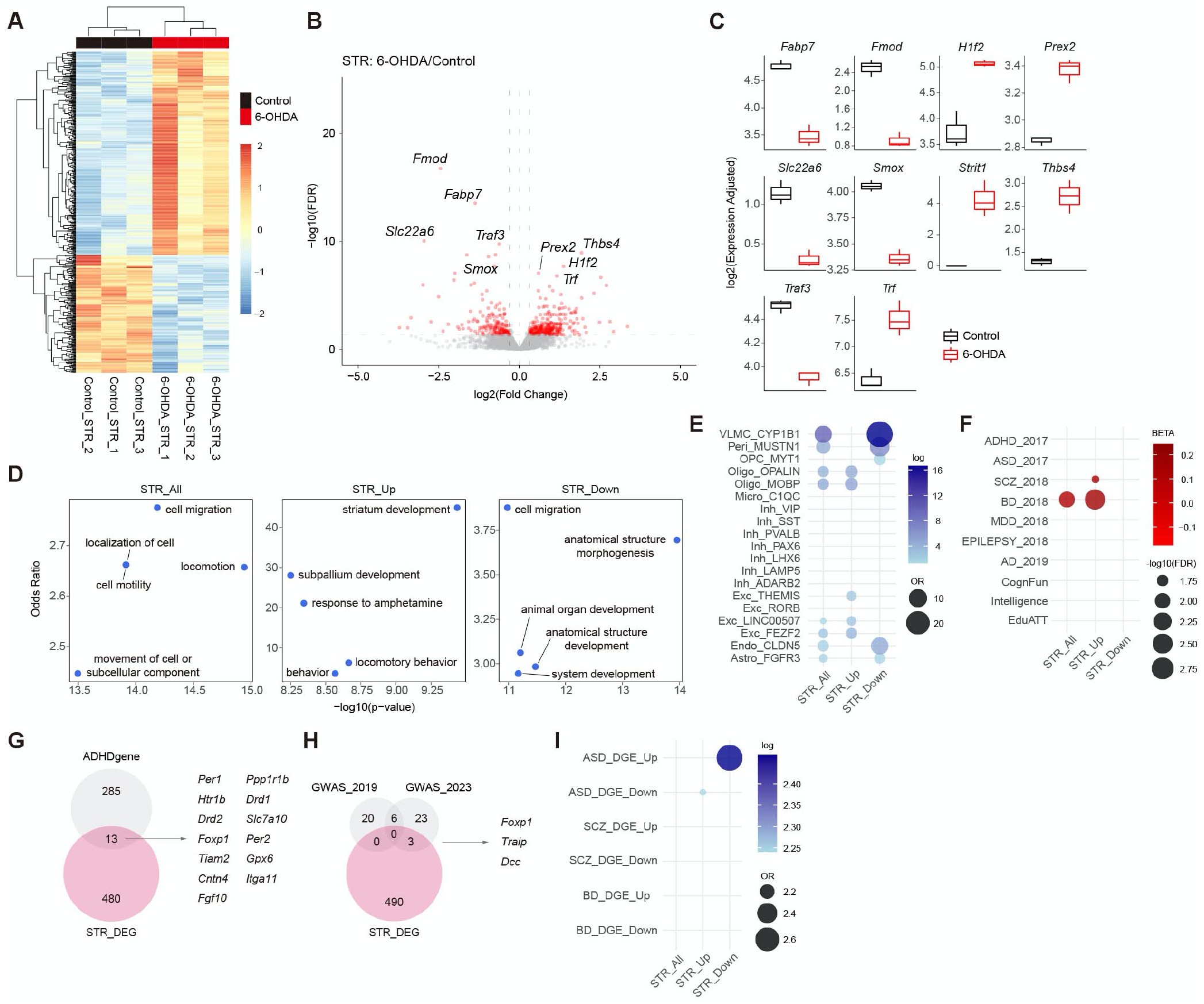
Transcriptomic remodeling in the STR of 6-OHDA-treated mice. (**A**) Heatmap of DEGs in the mouse STR at P24. (**B**) Volcano plot of STR-DEGs. (**C**) Bar plots showing the top 10 STR-DEGs. (**D**) GOs analysis of BP terms for STR-DEGs in all, upregulated, and downregulated genes, respectively. (**E**) Dot plots showing enrichment of STR-DEGs across cell types. (**F**) Dot plots showing MAGMA beta values. (**G**) Venn diagram showing overlap between STR-DEGs and ADHDgene. Representative overlapping genes, including *DRD2* and *FOXP1*, are highlighted. (**H**) Venn diagram showing overlap between STR-DEGs and GWAS-associated genes. Representative overlapping genes, including *FOXP1, DCC*, and *FOXP2*, are highlighted. (**I**) Dot plots showing enrichment of STR-DEGs in disorder-specific human transcriptomic datasets.

Cell type enrichment analysis revealed that STR DEGs were enriched in VLMCs, pericytes, endothelial cells, excitatory neurons, oligodendrocytes, and astrocytes (Figure 3E). GWAS enrichment analysis showed enrichment for genes associated with SCZ and bipolar disorder (BD) (Figure 3F). In addition, 13 STR DEGs (*Per1, Htr1b, Drd2, Foxp1, Tiam2, Cntn4, Fgf10, Pppr1b, Drd1, Slc7a10, Per2, Gpx6*, and *Itga11*) overlapped with ADHDgenes^20^ (Figure 3G), and 9 STR DEGs (*Kdm4a, Ptprf, Foxp2, Sorcs3, Dusp6, Poc1b, Foxp1, Traip*, and *Dcc*) overlapped with two GWAS datasets^5,6^ (Figure 3H). STR DEGs were also enriched among both upregulated and downregulated human DEGs associated with ASD (Figure 3I). These findings indicate that early postnatal dopaminergic disturbance induces distinct transcriptomic remodeling in the STR, particularly in molecular programs related to development, locomotor regulation, and extracellular and non-neuronal processes.

### Early postnatal dopaminergic disturbance reshapes cortico-striatal co-expression networks

To identify groups of highly co-expressed genes altered by early postnatal dopaminergic disturbance, we performed WGCNA on bulk RNA-seq data from the PFC and STR and identified 32 modules (Figure 4A, Table S5, and Figure S4, S5). These modules were highly enriched in PFC and STR DEGs (Figure 4B). The midnightblue and red modules were positively correlated with 6-OHDA treatment, whereas the black, brown, and greenyellow modules were negatively correlated (Figure 4B). The darkgrey, darkred, grey60, lightgreen, and purple modules were preferentially associated with the PFC, whereas the darkorange, magenta, and white modules were preferentially associated with the STR (Figure 4B).

**Figure 4.**
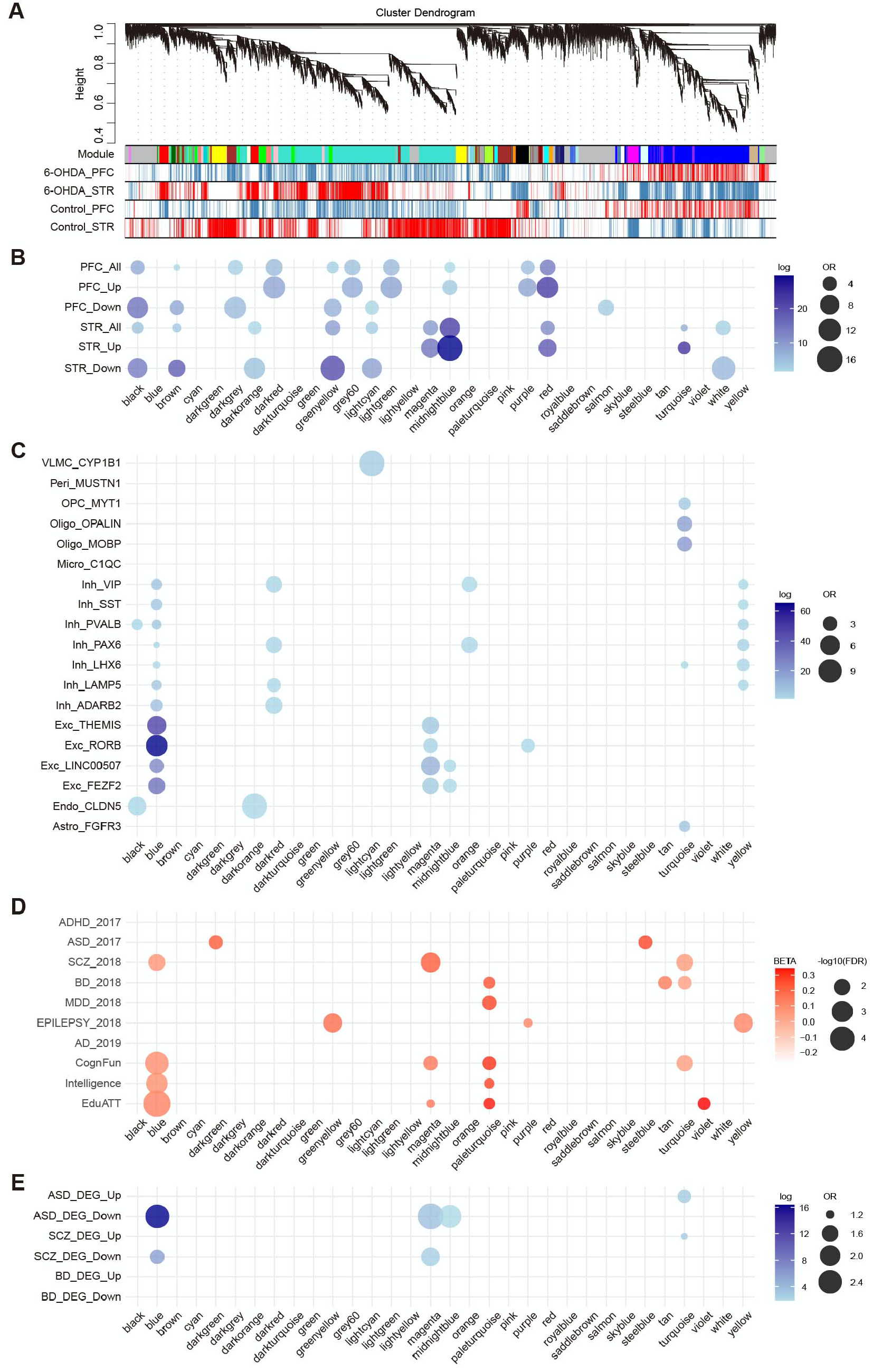
Gene co-expression networks in the PFC and STR of 6-OHDA-treated mice. (**A**) Co-expression network dendrogram of PFC and STR transcriptomes generated by weighted gene co-expression network analysis (WGCNA). (**B**) Dot plots showing enrichment of modules for DEGs. (**C**) Dot plots showing enrichment of modules across cell types. (**D**) Dot plots showing MAGMA beta values. (**E**) Dot plots showing enrichment of modules in disorder-specific human transcriptomic datasets.

Cell type enrichment analysis showed that the black and darkorange modules were enriched in endothelial cells, the blue module in inhibitory and excitatory neurons, the lightcyan module in VLMCs, the magenta, midnightblue, and purple modules in excitatory neurons, the darkred, orange, and yellow modules in inhibitory neurons, and the turquoise module in oligodendrocytes and astrocytes (Figure 4C). In GWAS enrichment analysis, no modules were specifically enriched in ADHD GWAS-risk loci (Figure 4D). However, the darkgreen and steelblue modules were enriched in ASD GWAS-risk loci, the blue, magenta, and turquoise modules for SCZ loci, and the paleturquoise, tan, and turquoise modules for BD loci (Figure 4D). The paleturquoise module was also enriched for major depressive disorder (MDD) loci, while the greenyellow, purple, and yellow modules were enriched for epilepsy loci. In addition, the blue, magenta, paleturquoise, turquoise, and violet modules were enriched for genes associated with cognitive function, intelligence, and educational attainment (Figure 4D).

In disorder-specific human transcriptome enrichment analysis, the blue, magenta, and turquoise modules were enriched in genes whose expression is altered in ASD and SCZ, while the midnightblue module was enriched in ASD genes (Figure 4E). These results indicate that early postnatal dopaminergic disturbance reshapes cortico-striatal co-expression networks linked to distinct cell types and neurodevelopmental, neuropsychiatric, and cognitive gene sets.

### Functional organization of region-biased co-expression modules

To investigate the biological functions represented by 6-OHDA-associated, PFC-specific, and STR-specific modules, we performed GO analysis for each module. The 6-OHDA-associated modules included the black, brown, greenyellow, midnightblue, and red modules (Figure 5A, and Table S6). The black module was associated with mitochondrial ATP synthesis coupled with electron transport, ATP synthesis coupled with electron transport, and cellular respiration. The brown module was associated with the tricarboxylic acid cycle, mitochondrial electron transport, ubiquinol to cytochrome c, and proteasomal ubiquitin-independent protein catabolic process. The greenyellow module was associated with nervous system development, generation of neurons, and biological regulation. The midnightblue module was associated with oxidative phosphorylation, response to reactive oxygen species, and response to hydrogen peroxide. The red module was associated with cilium organization, regulation of RNA metabolic process, ensheathment of neurons.

**Figure 5.**
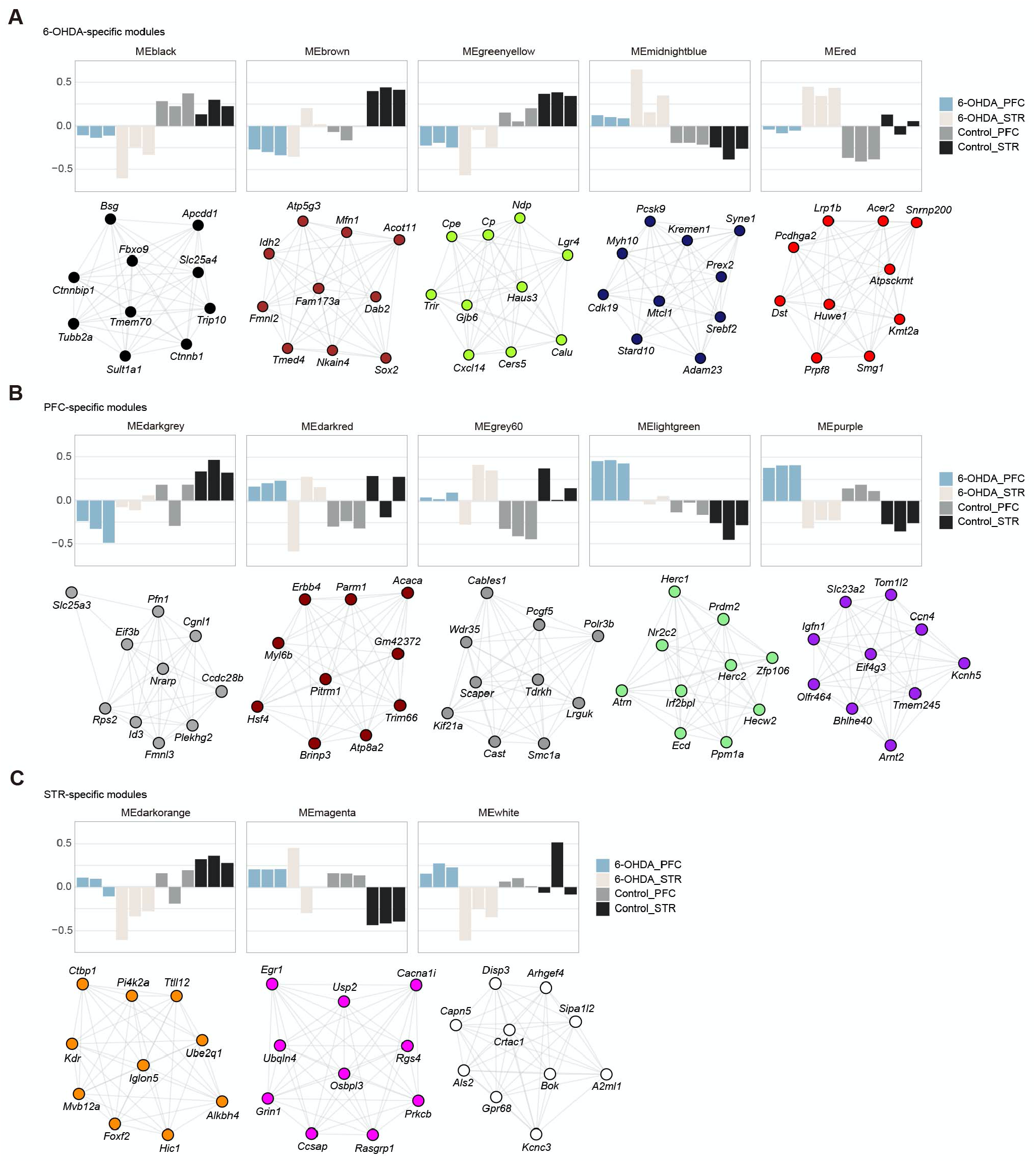
Co-expression networks of individual modules in 6-OHDA-treated mice. **(A–C)** Eigengene plots and co-expression networks of 6-OHDA-associated **(A)**, PFC-associated **(B)**, and STR-associated **(C)** modules, showing the top 10 hub genes. Hub genes play central roles in each co-expression network.

The PFC-specific modules included the darkgrey, darkred, grey60, lightgreen, and purple modules (Figure 5B, and Table S6). The darkgrey module was associated with cerebellum development, positive regulation of female receptivity, and response to thyroglobulin triiodothyronine. The darkred module was associated with trans-synaptic signaling, chemical synaptic transmission, and anterograde trans-synaptic signaling. The grey60 module was associated with cell-cell signaling, synaptic signaling, and trans-synaptic signaling. The lightgreen module was associated with phosphatidylinositol acyl-chain remodeling, regulation of cardiac conduction, and cell-cell adhesion via plasma-membrane adhesion molecules. The purple module was associated with saturated monocarboxylic acid metabolic process, unsaturated monocarboxylic acid metabolic process, and corticospinal tract morphogenesis.

The STR-specific modules included the darkorange, magenta, and white modules (Figure 5C, and Table S6). The darkorange module was associated with nitric oxide mediated signal transduction, response to macrophage colony-stimulating factor, and actin filament-based process. The magenta module was associated with response to lipoteichoic acid, RNA processing, and ncRNA transcription. The white module was associated with motile cilium assembly, positive regulation of leukocyte mediated immunity, and positive regulation of isotype switching to IgA isotypes. Together, these results indicate that early postnatal dopaminergic disturbance alters co-expression modules linked to developmental, synaptic, metabolic, immune-related, and transcriptional pathways in cortico-striatal regions.

## Discussion

In this study, we examined how early postnatal dopaminergic disturbance alters transcriptomic programs in the prefrontal cortex (PFC) and striatum (STR). Bulk RNA-seq identified 369 DEGs in the PFC and 493 DEGs in the STR, together with region-biased co-expression modules revealed by WGCNA. These findings indicate that early dopaminergic disturbance induces distinct molecular alterations across cortico-striatal regions during postnatal development. This interpretation is consistent with previous evidence implicating dopaminergic dysfunction in ADHD and related behavioral phenotypes^1,2^. It is also consistent with studies reporting frontostriatal circuit abnormalities in ADHD^5-7^. Our results extend these observations by showing that early dopaminergic perturbation is associated with region-specific remodeling of developmental, synaptic, metabolic, immune-related, and transcriptional programs in the PFC and STR.

A major finding of this study is the marked regional divergence between the PFC and STR following early postnatal dopaminergic disturbance. In the PFC, DEGs were enriched for anatomical structure development, regulation of developmental processes, plasma membrane signaling, and transcriptional regulation, suggesting that dopaminergic perturbation affects cortical developmental and signaling programs. In contrast, STR DEGs were enriched for striatal development, locomotion, extracellular matrix-related processes, and amphetamine response, indicating distinct molecular consequences for striatal maturation and functional regulation. These regional differences are notable in light of previous imaging studies implicating frontostriatal circuitry in ADHD^1,2^ and human transcriptomic studies reporting molecular alterations in ADHD^5-7,21^. Although a subset of DEGs was shared between the two regions, most changes were region-specific, supporting the idea that developmental dopaminergic disturbance reshapes local molecular programs according to regional context rather than producing a uniform brain-wide response.

Another notable feature of our dataset is that transcriptomic alterations extended beyond neuronal signatures. Cell-type enrichment analyses identified signals not only in excitatory and inhibitory neurons but also in VLMCs, pericytes, oligodendrocytes, and astrocytes. These findings suggest that early dopaminergic disturbance affects multiple cellular components of the developing brain. Oligodendrocytes are of particular interest because they form myelin sheaths around axons and provide metabolic support to neurons. Consistent with this, white matter abnormalities and altered connectivity have repeatedly been reported in ADHD^9,22,23^, and our previous comparative transcriptomic work also suggested that oligodendrocyte-related gene networks may be relevant to ADHD^24^. Astrocytes are likewise important regulators of extracellular neurotransmitter homeostasis, and striatal astrocytes express transporters and receptors implicated in catecholaminergic signaling^25,26^. In addition, VLMCs and pericytes may point to vascular- and barrier-associated responses. The WGCNA results further support this interpretation, as region-biased modules were linked to synaptic signaling, oxidative phosphorylation, mitochondrial function, RNA metabolism, actin-related processes, and immune-associated pathways. Together, these findings indicate that early dopaminergic disturbance reshapes cortico-striatal molecular organization at both the single-gene and network levels.

Our results also suggest that early dopaminergic disturbance may influence excitatory and inhibitory signaling, particularly in the STR. DEGs in the PFC were enriched in both excitatory and inhibitory neurons, whereas DEGs in the STR were enriched predominantly in excitatory neuronal signatures. Several region-biased modules were also associated with neuronal subtypes and synaptic signaling pathways. These observations are consistent with previous reports of reduced GABA concentrations in the motor cortex, striatum, and thalamus and elevated glutamate levels in frontal cortical regions in children with ADHD^27-31^. Although our transcriptomic data do not directly measure neurotransmitter levels or synaptic activity, they support the possibility that developmental dopaminergic disturbance alters molecular programs related to excitatory-inhibitory regulation in cortico-striatal systems.

External enrichment analyses showed overlap between DEG and module sets and genes implicated in ADHD and other neuropsychiatric disorders, including ASD, SCZ, and BD. These associations support the relevance of our dataset to neurodevelopmental and behavioral phenotypes and are broadly consistent with previous animal and human transcriptomic studies^21,32,33^. At the same time, these overlaps should be interpreted cautiously. The present study was designed to define the molecular consequences of early postnatal dopaminergic disturbance rather than to establish a comprehensive molecular signature of ADHD itself. Likewise, although 6-OHDA-treated mice displayed hyperactivity- and impulsivity-like phenotypes, the present findings are best interpreted as revealing developmental molecular programs associated with dopaminergic perturbation in cortico-striatal regions. This study is also limited by the use of a single pharmacological model and a single developmental time point. Future studies across additional models and developmental stages will be needed to determine which molecular changes are shared, persistent, or behaviorally causal. Nevertheless, our analyses identify region-specific transcriptomic and network-level changes in the PFC and STR and provide a useful framework for understanding how early dopaminergic disturbance may contribute to later behavioral phenotypes.

### Limitations of the study

This study has several limitations. First, our findings are based on a single pharmacological model of early postnatal dopaminergic disturbance. Second, the analyses were performed at a single developmental time point and therefore do not capture the temporal dynamics of transcriptomic changes across development. Third, although our data identify region-specific transcriptomic and network-level alterations in the PFC and STR, they do not directly establish which of these molecular changes are causally linked to later behavioral phenotypes. Future studies using additional models, developmental stages, and functional validation will be needed to determine which molecular programs are shared, persistent, and behaviorally relevant. Nevertheless, the present dataset provides a useful framework for understanding region-specific molecular responses to early dopaminergic disturbance during brain development.

## Resource availability

### Data availability

The data supporting the findings of this study are available from the corresponding author upon reasonable request. The NCBI Gene Expression Omnibus (GEO) accession number for the RNA-seq data are available at GSE277931.

### Code availability

Custom R codes and data to support the analysis, visualizations, functional, and gene set enrichments are available at https://github.com/BertoLabMUSC/Usui_ADHDproject/.

## Acknowledgements

We thank Miyoko Ieki, Shogo Togawa, and Koh Shinoda for support. This work was supported by the Japan Science and Technology Agency (JST) Center of Innovation Program (COI Program) (JPMJCE1310) to N.U. and S.S.; the Japan Society for the Promotion of Science (JSPS) Grant-in-Aid for Early-Career Scientists (23K14443, 26K18544) to M.D.; Foundation of Kinoshita Memorial Enterprise to M.D.; Hirose Foundation to M.D.; Niigata University Tsukada Medical Scholarship Fund to M.D.; Osaka Medical Research Foundation for Intractable Diseases to M.D. and N.U.; CNDD Genomics and Bioinformatics Core at MUSC (P20GM148302) to S.B.; JSPS Grant-in-Aid for Scientific Research (B) (23K27528) to N.U.; JSPS Grant-in-Aid for Scientific Research (C) (20K06872) to N.U.; JSPS Grant-in-Aid for Challenging Research (Exploratory) (24K22230) N.U.; Takeda Science Foundation to N.U.; Naito Foundation to N.U.; Mochida Memorial Foundation for Medical and Pharmaceutical Research to N.U.; Inamori Foundation to N.U.; SENSHIN Medical Research Foundation to N.U.; Shionogi infectious disease research promotion foundation to N.U.

## Author Contributions

**Miyuki Doi**: Conceptualization, Validation, Investigation, Writing - Original Draft, Writing - Review & Editing, Visualization, Funding acquisition. **Stefano Berto**: Methodology, Software, Writing - Review & Editing, Validation, Formal analysis, Investigation, Resources, Data Curation, Visualization, Funding acquisition. **Shoichi Shimada**: Writing - Review & Editing, Supervision, Funding acquisition. **Noriyoshi Usui**: Conceptualization, Methodology, Validation, Formal analysis, Investigation, Resources, Writing - Original Draft, Writing - Review & Editing, Visualization, Supervision, Project administration, Funding acquisition.

## Supplemental Information

Figures S1–S4, and Tables S1–S7.

## Methods

### Mice

All procedures were conducted in accordance with the ARRIVE guidelines and relevant official guidelines, approved by the Animal Research Committee of Niigata University (#SA01757) and The University of Osaka (#27-010). C57BL/6J (Japan SLC Inc., Shizuoka, Japan) male mice were used in this study, consistent with previous studies using this model. Mice were housed in cage (143 mm × 293 mm × 148 mm) in the barrier facilities of The University of Osaka under a 12 h light–dark cycle and given free access to water and food. Behavioral testing was performed between 10:00 and 16:00 h. Experimenters blinded to genotypes performed the test. The literature was referenced to convert the age of postnatal mice to human age^34^.

### 6-OHDA administration

A mouse model induced by early postnatal 6-OHDA treatment was generated as previously described^15,16^. This model exhibits ADHD-like behavioral phenotypes, including hyperactivity, impulsivity, and attention deficits^12^. Male mice were anesthetized by hypothermia on postnatal day (P) 5. Desipramine hydrochloride (20 mg/kg; #D3900, Merck, Darmstadt, Germany) was administered by unilateral subcutaneous injection, followed 30 min later by intracerebroventricular injection of saline or 6-OHDA hydrobromide containing 0.1% ascorbic acid as a stabilizer (25 μg dissolved in 3 μL saline; #H116, Merck). The intracerebroventricular injection was performed at a rate of 1.5 μL/min using a 10-μL Gastight Syringe (#1701RN, Merck) fitted with a 30G Small Hub RN Needle (#7803-07, 30GA, RN, 16 mm, 45°; Merck). Coordinates for injection were 0.6 mm lateral to the medial sagittal suture, 2.0 mm rostral to lambda, and 1.3 mm below the skin surface. After injection, pups were warmed on a heating pad at 37°C until recovery and then randomly returned to their dams. Pups remained with their dams until analysis at P24.

### Immunohistochemistry

Immunohistochemistry was performed as previously described^35^. Brains were fixed with 4% PFA in PBS overnight at 4°C, cryoprotected in 30% sucrose in PBS, then embedded in Tissue-Tek O.C.T. Compound (#4583, Sakura Finetek Japan Co.,Ltd., Osaka, Japan) for cryosectioning. Cryosections (20 μm thick) were placed in PBS. Sections were stained with the rabbit polyclonal anti-Tyrosine hydroxylase (1:500, #AB152, Merck, Burlington, MA) primary antibody. For fluorescence immunostaining, goat anti-rabbit IgG (H+L) cross-adsorbed secondary antibody conjugated to Alexa Fluor 488 (1:2,000, #A-11008, ThermoFisher, Waltham, MA) was applied, and cover glasses were mounted with Fluoromount/Plus (#K048, Diagnostic BioSystems, Pleasanton, CA). Images were collected using an all-in-one fluorescence microscope (BZ-X700, KEYENCE Corporation).

### Open field test

The open field test was performed as previously described^36^.Mice were placed in one of the corners of a novel chamber (W700 × D700 × H400 mm, #OF-36(M)SQ, Muromachi Kikai Co., Ltd., Tokyo, Japan) and were allowed to freely explore for 10 min. Locomotor activity was measured and tracked using ANY-maze behavior tracking software 7.33 (Stoelting Co., Wood Dale, IL, USA).

### Elevated plus maze test

Elevated plus maze test was performed as previously described^36^. Mice were placed in the center of the maze (open arms W54 × D297 mm; closed arms W60 × D300 × H150 mm; height from floor 400 mm, #EPM-04M, Muromachi Kikai Co., Ltd.) and allowed to explore freely the maze for 5 min. Time and distance were measured by ANY-maze (Stoelting Co.).

### Bulk RNA-seq

Bulk RNA-seq was performed using a service by Macrogen Japan (Kyoto, Japan) as previously described^36^. Total RNA was extracted from the PFC and STR of mice at P24 (N=3/condition) using the miRNeasy Mini Kit (#217004, Qiagen, Hilden, Germany). The PFC and STR were morphologically identified under a microscope after the brain was extracted, and each was carefully dissected and removed with tweezers. RNA integrity number (RIN) of total RNA was quantified by Agilent 2100 Bioanalyzer (Agilent Technologies) using Agilent RNA 6000 Pico Kit (#5067-1513, Agilent Technologies). Total RNA with RIN values of ≥8.9 were used for mRNA-seq library preparation. mRNA was purified from 500 ng total RNA by poly(A) method, and used for cDNA library preparation by TruSeq Stranded mRNA Library Prep (#20020594, Illumina, San Diego, CA, USA). The quality of cDNA Libraries were determined by 2100 Bioanalyzer using Agilent High Sensitivity DNA Kit (#5067-4626, Agilent Technologies). Libraries were sequenced as 101 bp paired-end reads on Illumina NovaSeq6000.

### RNA-seq data analyses

Reads were aligned to the mouse mm10 reference genome using STAR (v2.7.1a)^37^. For each sample, a BAM file was created containing mapped and unmapped reads across splice junctions. Secondary alignments and multi-mapped reads were removed using in-house scripts. Only uniquely mapped reads were retained for further analyses. Quality control metrics were assessed by the Picard tool (http://broadinstitute.github.io/picard/). Gencode annotation for mm10 (version M25) was used as reference alignment annotation and downstream quantification. Gene level expression was calculated using featureCounts (v2.0.1)^38^.

### Differential expression analysis

Counts were normalized using counts per million reads (CPM). Genes without reads in either sample were removed. Differential expression analysis was performed in R using DESeq2 (v1.34)^39^ with the following model: gene expression ∼ genotype. Log2 fold changes and p-values were estimated. P-values were adjusted for multiple comparisons using the Benjamini-Hochberg correction (FDR<0.05). Mouse Gene IDs were converted to Human Gene IDs using *biomaRt* package in R^40^.

### GO and enrichment analyses

Functional annotation of differentially expressed and co-expressed genes was performed using clusterProfiler (v4.2)^41^. Benjamini-Hochberg FDR (FDR<0.05) was applied as a multiple comparison adjustment. For cell-type specific enrichment of DEGs, scRNA-seq datasets from the mouse brain curated from PSY DEGS^42^ and scMouse Young (CZ CellXGene)^43^ were used. Gene set enrichment was performed in R using Fisher’s exact test with the following parameters: alternative = “greater”, confidence level = 0.95. We reported Odds Ratio (OR) and Benjamini-Hochberg adjusted p-values (FDR).

### WGCNA

WGCNA^44^ was performed using normalized RNA-seq data. The soft-threshold power was selected using the pickSoftThreshold function to approximate scale-free topology (R^2^>0.85), while also constraining mean connectivity to remain below 350 to avoid an overly dense network. This balance ensured both biological relevance and network interpretability. Modules were identified via dynamic tree-cutting with a deep split of 4, and Spearman’s rank correlation was used to compute module eigengene–treatment associations. Networks were constructed with blockwiseConsensusModules function with biweight midcorrelation (bicor). Power of 22 was selected for downstream analysis. We used these parameters for network construction: CorType = bicor, networkType = signed, TOMtype = signed, TOMDenom = mean, maxBlockSize = 16000, mergeCutHeight = 0.1, minCoreKME = 0.4, minKMEtoStay = 0.5, reassignThreshold = 1e-10, deepSplit = 4, detectCutHeight = 0.999, minModuleSize = 50.

### GWAS data and enrichment analyses

GWAS implementing MAGMA (v.1.07)^45^ was conducted. Protein coding genes included in the human gencode v.19 (19,346) were used as background for the gene-based association analysis. For each gene, SNPs located within the gene body (including exonic, intronic, and untranslated regions) as well as SNPs within 10 kb upstream and downstream were included in the gene annotation step. This mapping was performed using MAGMA’s --annotate function with default settings, using the NCBI build 37 (hg19) genome reference. The gene-based association tests incorporated linkage disequilibrium (LD) structure between SNPs, using the 1000 Genomes Project Phase 3 European reference panel. Gene P -values were computed by aggregating SNP-level summary statistics using the default SNP-wise mean model in MAGMA. Benjamini–Hochberg correction was applied, and significant enrichment is reported for FDR < 0.05. Summary statistics for GWAS of neuropsychiatric disorders and non-brain disorders were downloaded from the Psychiatric Genomics Consortium and other resources^5,46-59^.

### Statistical analysis

Bioinformatics statistics (Benjamini-Hochberg and Spearman’s rank correlation) and programs were performed using R as detailed in the corresponding sections above. Biological data are represented as means of biological independent experiments with ±standard error of the mean (SEM). Statistical analyses (unpaired *t*-test and Mann-Whitney test) were performed using Prism 9 (GraphPad Software, Boston, MA, USA). The numbers of samples and used statistics were described in each corresponding figure legend. Asterisks indicate p-values (***P<0.001, **P<0.01, *P<0.05). P<0.05 was considered to indicate statistical significance.

