## Supplementary Information for "Region-specific cortico-striatal transcriptomic remodeling following early postnatal dopaminergic disturbance"

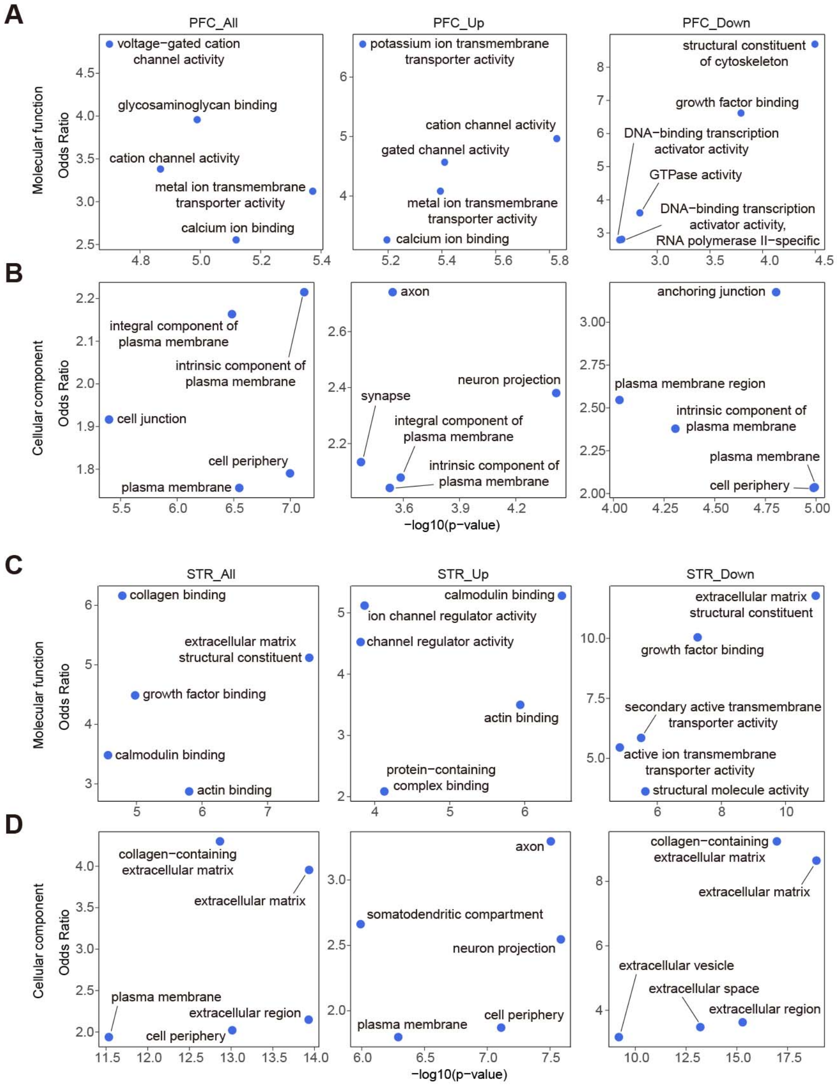


**Figure S1. Gene Ontology (GO) enrichment analysis of PFC- and STR-DEGs.**

(**A**, **B**) GOs terms for molecular function (**A**) and cellular component (**B**) of PFC-DEGs in all, upregulated, or downregulated genes, respectively.

(**C**, **D**) GOs terms for molecular function (**C**) and cellular component (**D**) of STR-DEGs in all, upregulated, or downregulated genes, respectively.

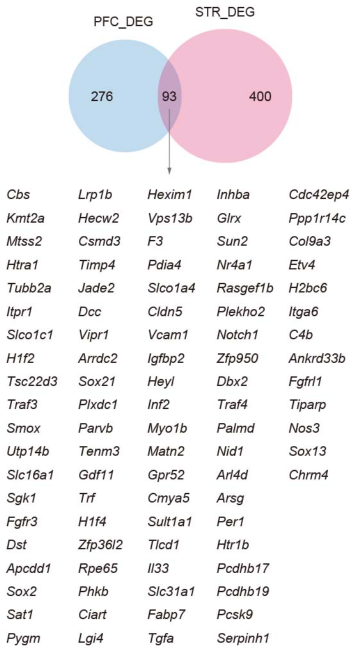


**Figure S2. Overlap between PFC- and STR-DEGs.**

Venn diagram showing the overlap between PFC-DEGs and STR-DEGs. A total of 93 DEGs were shared between the two regions.

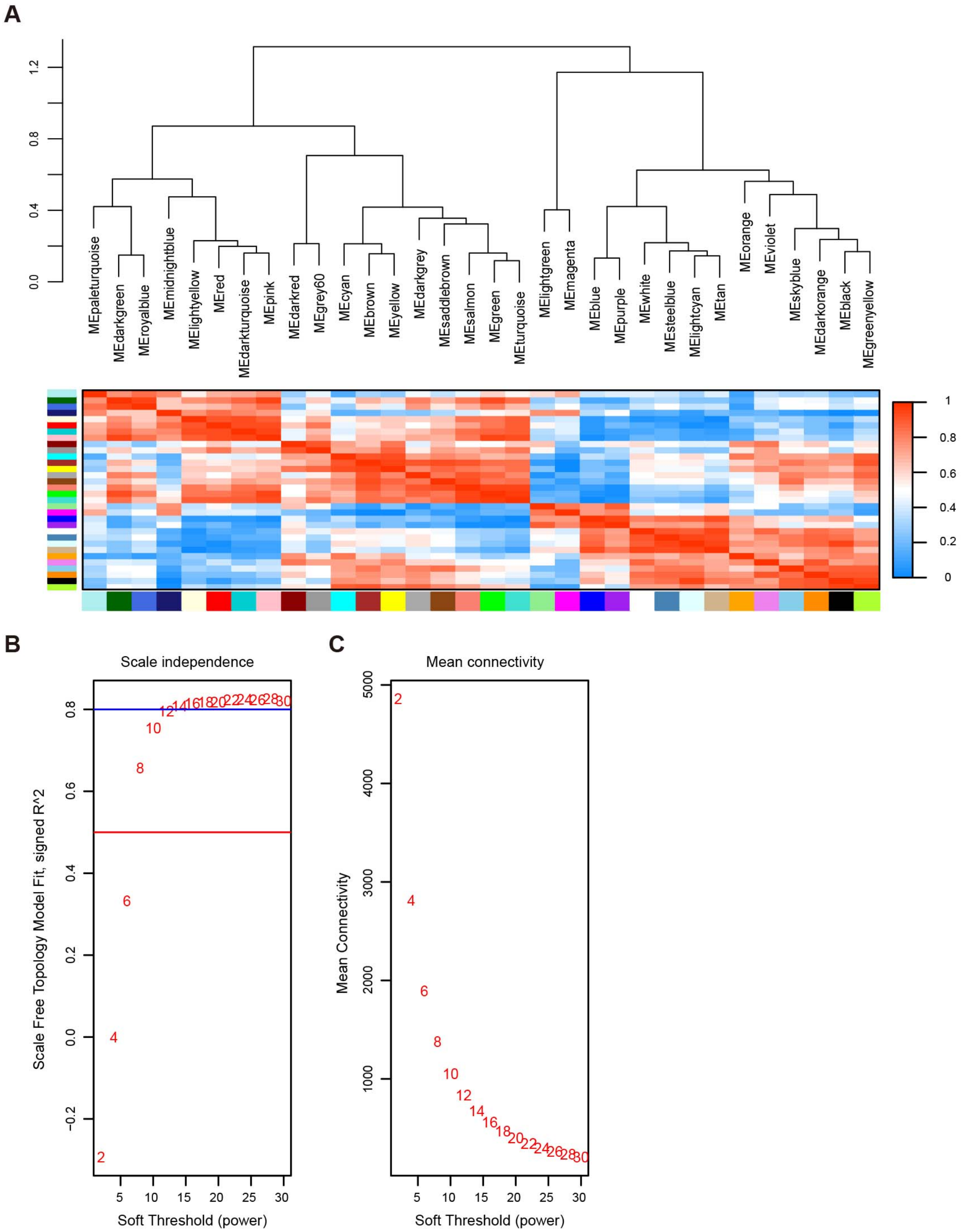


**Figure S3. Network topology and soft-threshold selection in WGCNA.**

(**A**) Eigengene network and heatmap.

(**B**) Scale-free topology fit index as a function of soft-threshold power.

(**C**) Mean connectivity as a function of soft-threshold power.

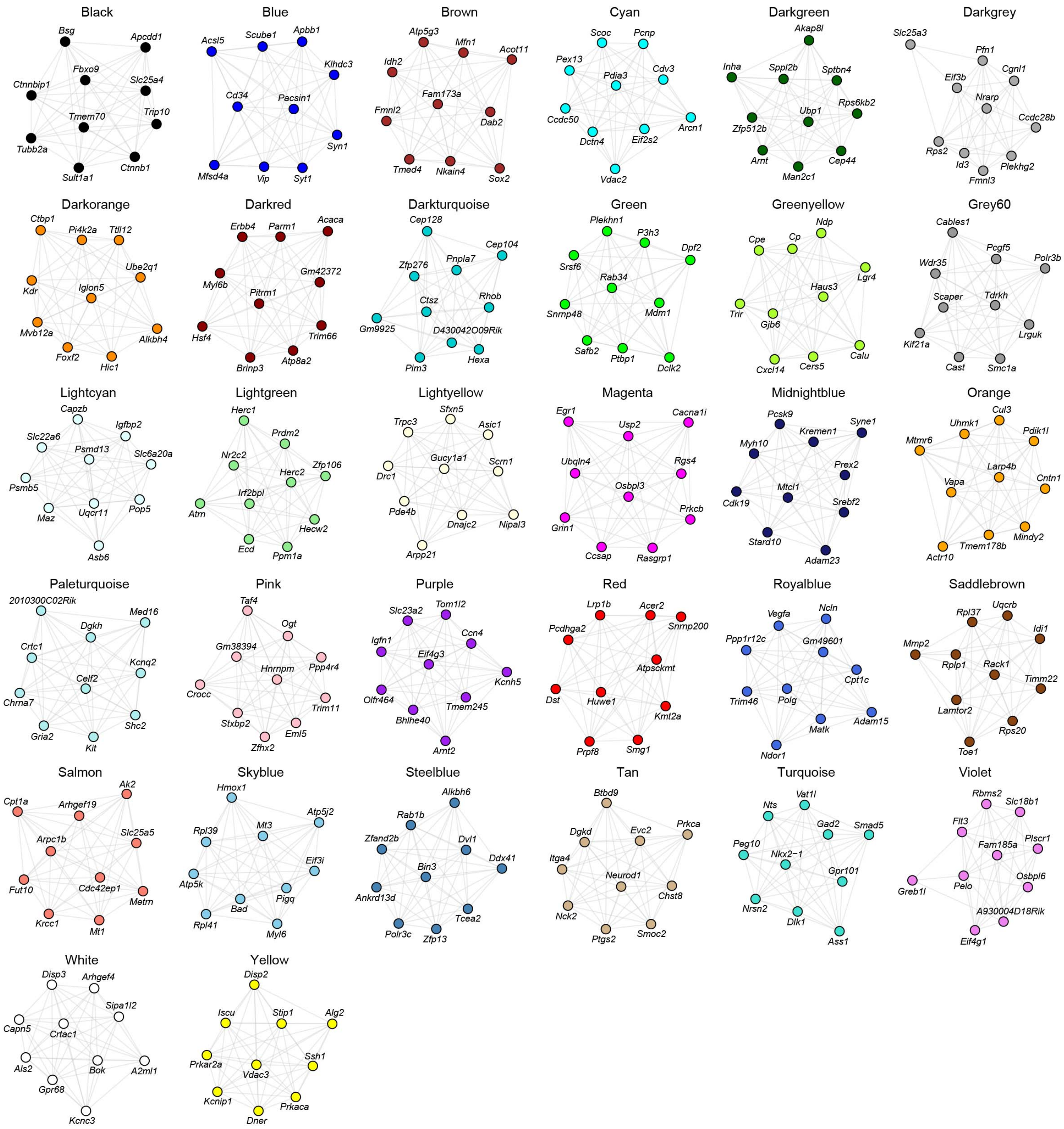


**Figure S4. Co-expression networks of individual WGCNA modules.**
Each module shows the co-expression network with the top 10 hub genes.

**Supplementary Tables**

Table S1. RNA-seq DEGs in the PFC of 6-OHDA-treated mice.
Table S2. GO analysis of PFC DEGs.
Table S3. RNA-seq DEGs in the STR of 6-OHDA-treated mice.
Table S4. GO analysis analysis of STR DEGs.
Table S5. WGCNA module outputs in 6-OHDA-treated mice.
Table S6. GO analysis of WGCNA modules.
Table S7. Overlap between WGCNA module genes and ADHDgenes.
